# Leveraging Tools from Autonomous Navigation for Rapid, Robust Neuron Connectivity

**DOI:** 10.1101/2020.04.30.070755

**Authors:** Nathan Drenkow, Justin Joyce, Jordan Matelsky, Reem Larabi, Jennifer Heiko, Dean Kleissas, Brock Wester, Erik C. Johnson, William Gray-Roncal

## Abstract

As biological imaging datasets continue to grow in size, extracting information from large image volumes presents a computationally intensive challenge. State-of-the-art algorithms are almost entirely dominated by the use of convolutional neural network approaches that may be difficult to run at scale given schedule, cost, and resource limitations. We demonstrate a novel solution for high-resolution electron microscopy brain image volumes that permits the identification of individual neurons and synapses. Instead of conventional approaches whereby voxels are labelled according to the neuron or neuron segment to which they belong, we instead focus on extracting the underlying brain graph represented by synaptic connections between individual neurons while also identifying key features like skeleton similarity and path length. This graph represents a critical step and scaffold for understanding the structure of neuronal circuitry. Our approach recasts the segmentation problem to one of path-finding between keypoints (i.e., connectivity) in an information sharing framework using virtual agents. We create a family of sensors which follow local decision-making rules that perform computationally cheap operations on potential fields to perform tasks such as avoiding cell membranes and finding synapses. These enable a swarm of virtual agents to efficiently and robustly traverse three-dimensional datasets, create a sparse segmentation of pathways, and capture connectivity information. We achieve results that meet or exceed state-of-the-art performance at a substantially lower computational cost. This tool offers a categorically different approach to connectome estimation that can augment how we extract connectivity information at scale. Our method is generalizable and may be extended to biomedical imaging problems such as tracing the bronchial trees in lungs or road networks in natural images.

## 1 Introduction

Connectomics is the study of structural and functional connections in the nervous system. Recent advances have allowed for large-scale imaging at the nanoscale, providing new capabilities for identifying single cells and their connections within larger brain networks. However, accurately and rapidly estimating these connections is a major bottleneck. Our approach represents a different way to directly map the connectivity between all neurons in a volume, as well as targeted sparse tracing of single neuron. Current approaches often frame circuit reconstruction as a segmentation problem that aims to label every pixel according to the structure to which it belongs (e.g., neuron, synapse, membrane) [1, 2]. Such methods are not optimized for graph generation and typically only operate over local sub-volumes due to computational or memory limitations that arise as a result of the large size of neural datasets (e.g., terabytes or petabytes of raw data). Scaling these approaches often leads to accumulated errors (e.g., large-scale merges) and sensitivity to noise (e.g., poor quality or missing data, misalignments, ambiguities from imaging artifacts).

In this work we explore a new approach to automated tracing. The state-of-the-art neuron segmentation algorithm, flood-filling networks [3], is reported to take on the order of 4.6 PFLOPs (an immense amount of computation) for a 5200×5200×5120nm subvolume and illustrates how connectome reconstruction is currently beyond the capabilities of many research groups. Further, much of the information produced may be unnecessary to explore initial questions leveraging directed, weighted graphs between neurons (e.g., information pathways, typical and pathological circuit patterns).

Our proposed approach seeks to provide a fast, scalable, and CPU-parallelizable method for generating neuron connectivity traces in EM volumes as an alternative to current approaches that are more difficult to scale and are driven by expensive GPUs. Our method allows for a varying trade-off between speed and accuracy, producing coarse models that can be refined over time. This method provides both neuron topology as well as approximate morphology (sometimes referred to as a ‘sketch connectome,’) but also allows for direct generation of brain network connectivity.

Our technique is inspired by previous work on dynamic co-fields (DCF) [4, 5] which introduced a method for controlling the movement of a mobile autonomous agent swarm through the use of coordination fields. We re-frame the neuron tracing problem as one where autonomous agents trace an EM volume utilizing the outputs of a series of weighted potential fields as local decision making rules to seek out unexplored regions and search for synapses. Dynamic weighting of these fields allows our model to be robust to data defects, one of the current challenges in the field.

We demonstrate the fast, scalable and robust qualities of this agent-based approach for both large-scale connectome generation as well as targeted neuron tracing. We also demonstrate how brain graphs can be directly derived from agent traces. We believe that optimizing for topology instead of morphology provides a more rapid and complementary method for global scene understanding and network reconstruction, compared to current state-of-the-art methods.

## 2 Methods

### 2.1 Overview

Our approach recasts the pixel-level segmentation problem to one of real-time graph extraction using virtual agents. These agents are spawned at putative synapse locations and at regular intervals throughout the volume. At each time step, agents use characteristics of the surrounding 3D space combined with local decision making rules implemented as sensors, each of which contributes updates to the agent’s velocity (i.e., direction and speed) illustrated in Figure 1. Equipped with sensors, agents frequently and efficiently sample information about their own history and their local environment and modify their behavior according to new observations.

**Figure 1:**
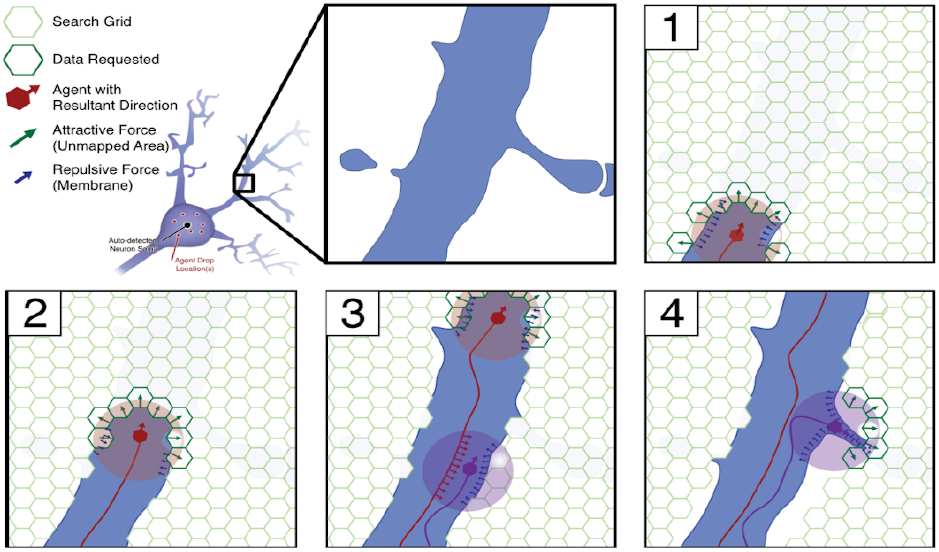
To trace neurons, image data are re-interpreted as a series of fields which contribute forces that motivate agent behavior. (1-3) Agents are repelled from locations that have already been traversed by other agents. (3) An agent is spawned and is simultaneously attracted to unexplored regions and repelled from membranes. (4) Cooperative exploration motivates exploration of small processes in search of synapses.

At the conclusion of an experiment, agents are merged through spatial proximity. These combined location histories result in point sets that each represent paths within a single neuron. These trajectories are used with the synapses encountered by the agents to trivially form the output graph (i.e., synaptic connections between neurons). The agent-based approach enables parallel cooperative tracing of multiple neurons. Agents can be trivially parallelized across CPU cores and can easily be run on consumer hardware.

### 2.2 Fields

In the DCF paradigm, coordination fields, *F*_*i*_, encode contextual information that agents sense and rely on to dictate their behavior. Fields provide a representation of the input space, and sensors perform arithmetic operations on these fields to yield useful decisions. In our EM connectomics application, relevant fields include membrane and synapse maps, agent exploration fields, and boundary maps.

Membrane fields are obtained by performing a cheap segmentation of the EM data (using a U-Net architecture) [6], applying a threshold to the probability maps, and performing a morphological closing operation to remove small gaps. A Sobel filter is then used to find the vectors pointing away from the edges, and these values are stored in a lookup table. By varying filter sizes, the sensor can be robust to the defects that result from imperfections that remain in the post-processed U-Net output. The synapse field is obtained by a similar approach, but with an opposite direction to attract agents. Sensors are tuned to minimize accidental membrane crossings, which result in false merges.

Another field type is used to manage boundary crossings. Agents that hit a data boundary can be respawned in an adjacent sub-volume, aiding in merging decisions and facilitating parallelization. This allows agents to scale to arbitrarily large image volumes without a costly and error-prone merging step that can adversely affect graph generation.

An exploration field is maintained by the swarm to account for visited regions; agents will sense locally and move towards unexplored areas. In a distributed framework, agents may broadcast and receive updates from other agents that can be used to update their local field and promote collaborative behavior.

### 2.3 Agents

Agents are virtual entities that are atomically defined by their real-valued position 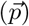 and velocity vector 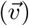 and are equipped with a set of sensors which allow them to observe their environment. At each time step, every agent modifies its velocity by the weighted sum of the unit vectors returned by each of its sensors and subsequently updates its position by this velocity. Agents can be throttled by constraining them to a maximum velocity (*s*_*max*_), thereby preventing runoff. Different agents may be given different sets of sensors, allowing some to be ‘specialists’ while others remain ‘generalists.’

### 2.4 Sensors

A sensor 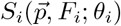 receives the current position and appropriate field *F*_*i*_ and is parameterized according to *θ*_*i*_ which may describe the range, direction, or other relevant aspects of the sensing operation. Sensors produce an influence vector 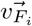 which summarizes the direction in which the agent should move next as determined by the local values of a particular field.

Sensors create influence vectors directly from field output, as is the case for the membrane avoidance sensor. Agents can also perform operations on these influence vectors to produce further useful results.

For example, taking the vectors orthogonal to the membrane avoidance sensor yields a useful wall follower sensor. Sensors can also perform operations on agents, like killing an agent when it hits a wall and spawning a new agent in a new unexplored area of the volume.

### 2.5 Agent-based framework

In our agent-based framework, described in Figure 2, we combine the co-fields, agents, and sensors as follows. Given a set of fields and sensors, our agent moves according to equation 1 whereby each sensor for each agent retrieves its corresponding influence vector *S*_*i*_(*p*_*t*_, *F*_*i*_; *θ*_*i*_) which is scaled by a weight *w*_*i*_. Each weighted vector is summed and added to the current agent velocity, and this new velocity is used to calculate the agents next position.

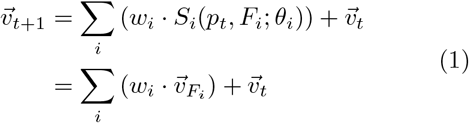

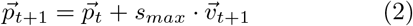

**Figure 2:**
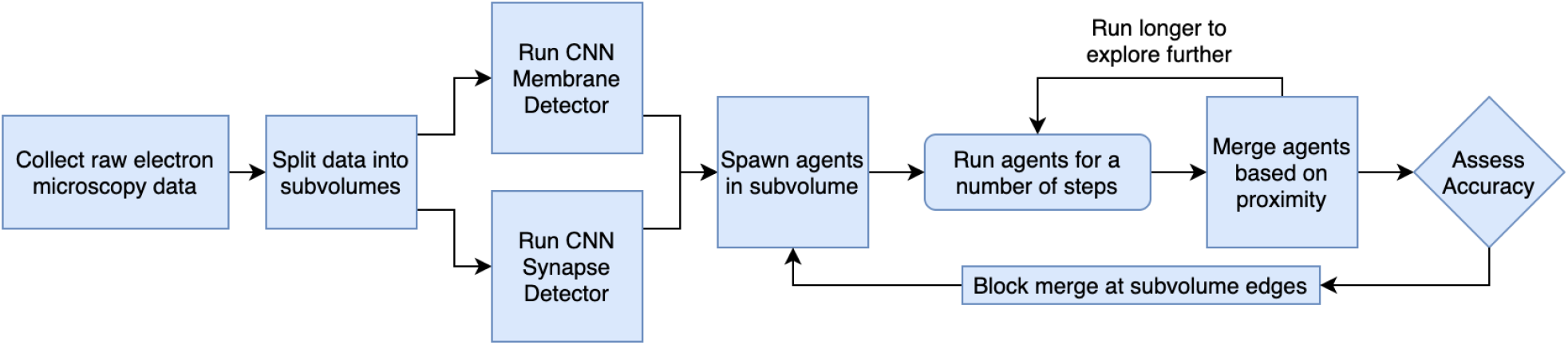
The agents workflow starts with an image volume, traces neuronal pathways with Agents, and creates output graphs.

As agents move through image space, agent state information is recorded including position and velocity histories as well as synapses visited, facilitating exploration. Agent consensus can be used to create confidence maps of neuron segmentation, which can be helpful for tracing coarse neuron morphologies even under suboptimal weight parameters.

### 2.6 Graph Generation

After exploring an image volume, the connectivity graph can be recovered by fusing agent path histories to determine synapse ownership and connectivity. We can fuse paths by considering voxel level overlap, and can extend these ideas by considering similar trajectories to resolve ambiguities. The outputs of the merge are the sets of agent path histories that are connected. By combining these merge sets with the identified synaptic connections, a graph can be created. When synapses are annotated as point annotations denoting the pre- and post-synaptic terminal locations (as is popular in fly data), the merged agents at the pre-synaptic site are connected to the merged agents at the post-synaptic site, creating a directed edge.

## 3 Experiments and Results

Our approach was primarily tested on the FIB-25 dataset [7]. This data is an approximately 27,000 cubic micron subvolume of a medullar column of a fruit fly (*Drosophila melanogaster*). This dataset has been used as a testbed for the development of segmentation algorithms such as state-of-the-art flood-filling networks (FFN) [3]. The volumes used in the current experiment were the published training and validation volumes from Janelia (250×250×250 and 520×520×520 voxels in size at a resolution of 8×8×8nm) [8]. As our comparison method, we used FFN’s pretrained weights from the authors[3]. FIB-25 has been manually segmented by expert researchers who have labelled every voxel of the dataset according to the unique neuron ID to which it belongs. Likewise, the synapses are labelled as pre-synaptic points connected to a series of post-synaptic locations. These together form the data used as ground truth, and these true synapses were used to compare agents and FFN path reconstructions.

Agent tracing runs were conducted using varying numbers of steps and numbers of agents. The results of 19 standardized runs are listed in figure 3 and show the relationship between increasing the number of agents and graph integrity.

**Figure 3:**
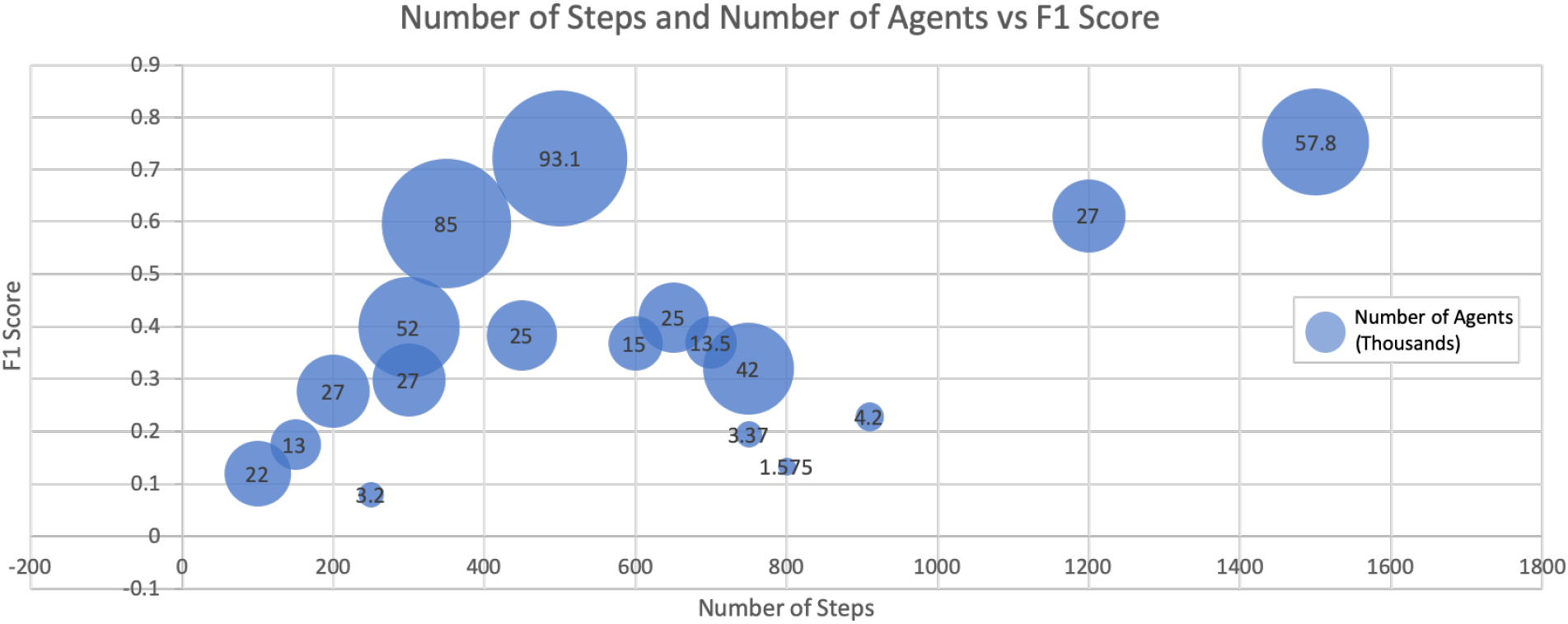
Increasing the number of agents and increasing the number of steps increase the graph accuracy of the merged run. We seek to minimize computation expense while ensuring good exploration and therefore neuron representation.

Graph integrity was measured based on neural reconstruction integrity (NRI) [9], a metric that measures graph similarity according to precision and recall of intracellular paths. Incorrect edges (additions or deletions) are penalized and used to create a precision and recall score reflective of graph quality. Additionally, an edge-based precision-recall metric was used that directly calculates scores based on incorrectly found (i.e., false positive) and omitted (i.e., false negative) edges. For this measurement, we construct line graphs [10]. This representation considers the synapses as nodes, and the edges as paths between synapses. These new graphs are built using the same pre- and post-synaptic synapse points, and reconstructions are compared to the ground truth line graph to determine precision and recall, with each edge representing a detection. These are reported in Table 1. Agents performs on par with state-of-the-art at a fraction of the computational cost and on hardware that more easily allows for large-scale parallelization. The tracing results reported on in the table are from a run with 1500 steps and about 50,000 agents spawned both preferentially at synapses as well as uniformly throughout the volume. In preliminary experiments, we have observed that the same weights perform well on other datasets, suggesting that extensive parameter tuning is not required for transfer.

**Table 1:** Results for experiments run on the FIB-25 dataset

|  | Graph |  | NRI |  |  | Edge |  |  |
| --- | --- | --- | --- | --- | --- | --- | --- | --- |
|  | nodes | edges | precision | recall | f1 | precision | recall | f1 |
| Ground Truth | 589 | 1137 | - | - | - | - | - | - |
| FFN | 779 | 1251 | 0.767 | .709 | .737 | .692 | .790 | .738 |
| Agents (ours) | 745 | 1198 | .824 | .693 | .753 | .753 | .741 | .746 |

In order to optimize sensor weights, we ran a grid search across the sensor parameter space using gradient descent with NRI as the cost function. Both FFN and our approach were trained and then validated on the aforementioned training and validation sets provided by Janelia. We aimed to tune our error profile to emphasize graph splits over merges since the workflows of many successful high-throughput connectomics projects, prefer to quickly stitch together path fragments rather than break up merges. This operating point is created by tuning down the wall following sensor and turning up the wall avoidance sensor, causing agents to segment more conservatively.

For the purposes of the above comparison, these graphs were generated from relatively small volumes. This algorithm can be readily adapted to batch processing frameworks (e.g., [11]) and large spatial databases (e.g., [12]).

## 4 Discussion and Conclusions

Agent-based path tracing is complementary to that of existing dense segmentation approaches and can provide crucial context for voxel-accurate dense measurements in targeted regions using a coarse-to-fine approach. We emphasize that this approach is different than the conventional methods used to estimate connectome graphs and offers an attractive alternative for rapidly, cheaply, and robustly creating connectivity maps. As datasets grow in size, processing these volumes can be a major challenge for research groups, and this technique allows users to focus less on perfect alignment and sample preparation and avoid some of the more common challenges associated with scalability. This method is also easily parallelizable across CPUs allowing computer clusters to segment a large volume very quickly and at low cost.

For some applications, detailed morphology is critical, and this technique will require post-processing to produce dense maps; although this has been addressed in other work [13]. We observe that Agents reaches many, but not all small processes; tortuous spine necks can be challenging as can be seen in Figure 4, as well processes that appear disconnected in adjacent slices. Future work will explore more powerful sensing strategies and post-processing (e.g., learning) approaches to address these scenarios, as well as transfer to other datasets. Our techniques can be extended naturally to other graph reconstruction challenges such as those in magnetic resonance imaging, CLARITY, array tomography and X-ray microtomography (XRM). We believe that this technique is synergistic with existing methods and will allow scientists to more rapidly understand connectivity in large, high-resolution datasets.

**Figure 4:**
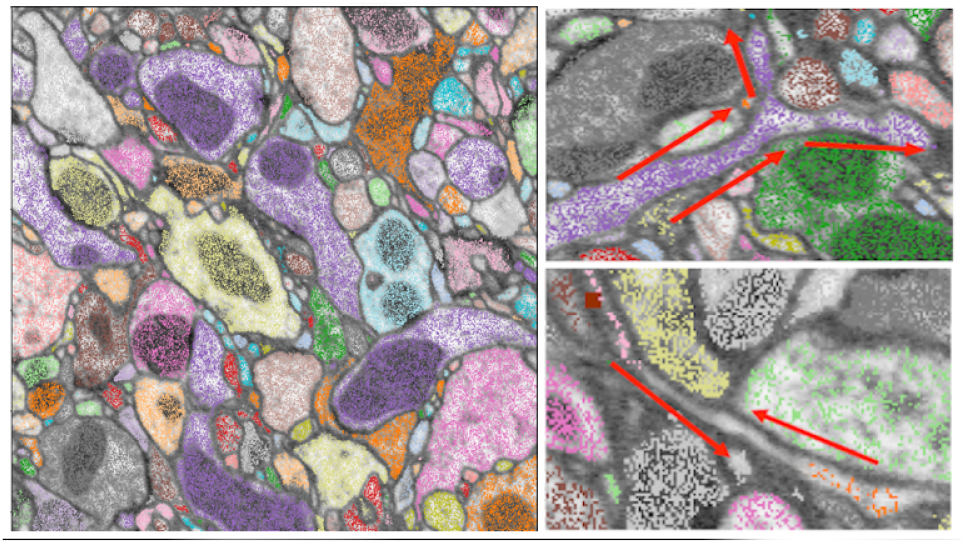
(Left) A slice of segmentation output, visualized as a 2D cross-section of the merged paths from an agent run. The background is the EM image slice representing the input data, and the dark areas within some of the boundaries are mitochondria, which can be distractors in creating an output graph. (Middle) Agents populate and move along branching passageways, efficiently tracing neuron morphologies (Right) The narrowest passages are difficult for agents to pass through, even with optimized sensor weights. The membrane boundary sensors push agents away from these passages. There is a trade-off between robustness to false gaps that arise as a result of mis-alignments or image processing artifacts and robustly traversing thin pathways. For many workflows, these errors are quick for human annotators to correct

## 5 Acknowledgements

This material is based upon work supported by the Office of the Director of National Intelligence (ODNI), Intelligence Advanced Research Projects Activity (IARPA), via IARPA Contract No. 2017-17032700004-005 under the MICrONS program. The views and conclusions contained herein are those of the authors and should not be interpreted as necessarily representing the official policies or endorsements, either expressed or implied, of the ODNI, IARPA, or the U.S. Government. The U.S. Government is authorized to reproduce and distribute reprints for Governmental purposes notwithstanding any copyright annotation therein. Research reported in this publication was also supported by the National Institute of Mental Health of the National Institutes of Health under Award Numbers R24MH114799 and R24MH114785. The content is solely the responsibility of the authors and does not necessarily represent the official views of the National Institutes of Health. This work was completed with the support of the CIRCUIT initiative circuitinstitute.org, and JHU/APL Internal Research Funding.

## References

[1] Parag, Toufiq and Tschopp, Fabian and Grisaitis, William and Turaga, Srinivas C and Zhang, Xuewen and Matejek, Brian and Kamentsky, Lee and Lichtman, Jeff W and Pfister, Hanspeter, “Anisotropic EM Segmentation by 3D Affinity Learning and Agglomeration,” arXiv preprint arXiv:1707.08935, 2017.

[2] Nunez-Iglesias, Juan and Kennedy, Ryan and Plaza, Stephen M and Chakraborty, Anirban and Katz, William T, “Graph-based active learning of agglomeration (GALA): a Python library to segment 2D and 3D neuroimages,” Frontiers in neuroinformatics, vol. 8, p. 34, 2014.

[3] Januszewski, Michał and Kornfeld, Jörgen and Li, Peter H and Pope, Art and Blakely, Tim and Lindsey, Larry and Maitin-Shepard, Jeremy and Tyka, Mike and Denk, Winfried and Jain, Viren, “High-precision automated reconstruction of neurons with flood-filling networks,” Nature methods, vol. 15, no. 8, pp. 605–610, 2018.

[4] Mamei, Marco and Zambonelli, Franco and Leonardi, Letizia, “Cofields: a physically inspired approach to motion coordination,” IEEE Pervasive Computing, vol. 3, no. 2, pp. 52–61, 2004.

[5] Chalmers, Robert and Scheidt, David and Neighoff, Todd and Witwicki, S and Bamberger, Robert, “Cooperating unmanned vehicles,” in AIAA 1st Intelligent Systems Technical Conference, p. 6252.

[6] Ronneberger, Olaf and Fischer, Philipp and Brox, Thomas, “U-net: Convolutional networks for biomedical image segmentation,” in International Conference on Medical image computing and computer-assisted intervention. Springer, 2015, pp. 234–241.

[7] Takemura, Shin-ya and Xu, C Shan and Lu, Zhiyuan and Rivlin, Patricia K and Parag, Toufiq and Olbris, Donald J and Plaza, Stephen and Zhao, Ting and Katz, William T and Umayam, Lowell and others, “Synaptic circuits and their variations within different columns in the visual system of Drosophila,” Proceedings of the National Academy of Sciences, vol. 112, no. 44, pp. 13 711–13 716, 2015.

[8] S. Plaza, “Examples for NeuroProof.” [Online]. Available: https://github.com/janelia-flyem/neuroproof_examples

[9] Reilly, Elizabeth P and Garretson, Jeffrey S and Gray Roncal, William R and Kleissas, Dean M and Wester, Brock A and Chevillet, Mark A and Roos, Matthew J, “Neural reconstruction integrity: A metric for assessing the connectivity accuracy of reconstructed neural networks,” Frontiers in neuroinformatics, vol. 12, p. 74, 2018.

[10] Gray Roncal, Williamł and Kleissas, Dean and Vogelstein, Joshua and Manavalan, Priya and Lillaney, Kunal and Pekala, Michael and Burns, Randal and Vogelstein, R. Jacob and Priebe, Carey and Chevillet, Mark and Hager, Gregory, “An automated images-to-graphs framework for high resolution connectomics,” Frontiers in Neuroinformatics, vol. 9, p. 20, 2015. [Online]. Available: https://www.frontiersin.org/article/10.3389/fninf.2015.00020

[11] “Toward A Reproducible, Scalable Framework for Processing Large Neuroimaging Datasets, url=http://dx.doi.org/10.1101/615161, doi=10.1101/615161, publisher=Cold Spring Harbor Laboratory, author=Johnson, Erik C. and Wilt, Miller and Rodriguez, Luis M. and Norman-Tenazas, Raphael and Rivera, Corban and Drenkow, Nathan and Kleissas, Dean and LaGrow, Theodore J. and Cowley, Hannah and Downs, Joseph and et al., year=2019, month=Apr, journal = arXiv.”

[12] D. Kleissas, R. Hider, D. Pryor, T. Gion, P. Manavalan, J. Matelsky, A. Baden, K. Lillaney, R. Burns, D. D’Angelo et al., “The block object storage service (bossDB): A cloud-native approach for petascale neuroscience discovery,” bioRxiv, p. 217745, 2017.

[13] Helmstaedter, Moritz and Briggman, Kevin L. and Turaga, Srinivas C. and Jain, Viren and Seung, H. Sebastian and Denk, Winfried, “Connectomic reconstruction of the inner plexiform layer in the mouse retina,” Nature, vol. 500, no. 7461, p. 168, 2013.

